# FLAb: Benchmarking deep learning methods for antibody fitness prediction

**DOI:** 10.1101/2024.01.13.575504

**Authors:** Michael Chungyoun, Jeffrey Ruffolo, Jeffrey Gray

## Abstract

The successful application of machine learning in therapeutic antibody design relies heavily on the ability of models to accurately represent the sequence-structure-function landscape, also known as the fitness landscape. Previous protein benchmarks (including The Critical Assessment of Function Annotation [33], Tasks Assessing Protein Embeddings [23], and FLIP [6]) examine fitness and mutational landscapes across many protein families, but they either exclude antibody data or use very little of it. In light of this, we present the Fitness Landscape for Antibodies (FLAb), the largest therapeutic antibody design benchmark to date. FLAb currently encompasses six properties of therapeutic antibodies: (1) expression, (2) thermostability, (3) immunogenicity, (4) aggregation, (5) polyreactivity, and (6) binding affinity. We use FLAb to assess the performance of various widely adopted, pretrained, deep learning models for proteins (IgLM [28], AntiBERTy [26], ProtGPT2 [11], ProGen2 [21], ProteinMPNN [7], and ESM-IF [13]); and compare them to physics-based Rosetta [1]. Overall, no models are able to correlate with all properties or across multiple datasets of similar properties, indicating that more work is needed in prediction of antibody fitness. Additionally, we elucidate how wild type origin, deep learning architecture, training data composition, parameter size, and evolutionary signal affect performance, and we identify which fitness landscapes are more readily captured by each protein model. To promote an expansion on therapeutic antibody design benchmarking, all FLAb data are freely accessible and open for additional contribution at https://github.com/Graylab/FLAb.

## 1 Introduction

The innate and adaptive immune systems are pivotal for safeguarding the human body, with antibodies acting as specialized proteins evolved to combat diseases. Antibody engineering exploits their therapeutic potential, resulting in over 150 therapeutic antibodies targeting diverse diseases [4]. The efficacy of therapeutic antibody candidates hinges on achieving a delicate balance of drug-like biophysical properties, often characterized by intricate trade-offs where enhancing one property may compromise others [19].

The flourishing field of AI now shows promise in driving antibody design by generating new and diverse therapeutic candidates that have desirable biophysical characteristics in significantly less time. [5]. As the diversity of deep learning approaches increases [10, 29, 32, 31, 9, 18, 2, 3], it becomes vital to converge on a systematic benchmark for evaluating performance. Current antibody design methods are evaluated with less informative metrics (like native sequence recovery), which does not does provide a clear indication of therapeutic potential. In this study, we curate experimental fitness data from eight studies spanning antibody expression, thermostability, immunogenicity, aggregation, polyreactivity, and binding affinity into the Fitness LAndscape for Antibodies (FLAb). Then, we assess a collection of models relevant to antibodies for their ability to correlate likelihoods to fitness properties. Our long term vision is that FLAb will help the development of models that can filter new antibody design candidates more efficiently than what is more typically done experimentally.

## 2 Related work

Previous endeavors to establish benchmarks for function prediction have laid a foundation for protein engineers to assess new designs. The Critical Assessment of Function Annotation (CAFA) aims to assign gene ontology classes to proteins [33]. The Task Assessing Protein Embeddings (TAPE) evaluates different pretrained models in predicting three protein structure properties (remote homology, secondary structure, residue contacts), as well as two fitness properties (fluorescence and stability) [23]. Dallago *et al*. introduced FLIP, which examines complex fitness landscapes and performance across a diverse set of proteins encompassing various functions [6]. However, these benchmarks exclude antibody data, motivating us to curate publicly accessible antibody fitness data. Related work has also assembled antibody sequence and structure data, notably the Observed Antibody Space (OAS [22]) of annotated sequences from immune repertoires and the Structural Antibody Database (SAbDab [8]) of all antibody structures available in the Protein Data Bank. These databases focus on sequence and structure, but not fitness metrics.

## 3 Results

### 3.1 Fitness Landscape Collection

Jain *et al*. define the characteristics that comprise antibody developability, which includes (1) high-level of expression, (2) high conformational and colloidal stability, (3) low immunogenicity, (4) high binding affinity towards the target antigen, (5) a low propensity for aggregation, and (6) low polyreactivity [16]. To assess the efficacy of protein design models in capturing essential characteristics of therapeutic antibodies, we have compiled a collection (Table 1) of 17 mutational landscapes of distinct antibody families with a total of 13,384 associated fitness metrics relevant to Jain *et al*.’s definition of antibody developability [12, 17, 20, 24, 27, 30, 16, 15]. Each sequence is associated with at least one fitness label pertaining to the six aforementioned developability factors. Additional detail on fitness landscape descriptions and datasets collected can be found in Supp. A.2. A glossary of domain specific terminology is provided in Supp. A.13. We hypothesize that if a protein model displays statistically significant correlations with the antibody fitness landscapes, they can be considered reliable predictors for new therapeutic antibody design candidates.

**Table 1.** Number of unique fitness values from available antibody datasets.

| Antibody set | Exp.<br>($\mu\text{g/mL}$) | $T_m$<br>($^{\circ}\text{C}$) | Imm.<br>(% ADA) | Binding<br>(nM) | Agg.<br>(Wv shift) | Poly.<br>(min) |
| --- | --- | --- | --- | --- | --- | --- |
| GSK CA1 | 34 | 34 | - | 29 | - | - |
| GSK CA2 | 25 | 22 | - | 22 | - | - |
| GSK CA3 | 11 | 8 | - | 11 | - | - |
| GSK CA4 | 24 | 24 | - | 19 | - | - |
| Hie C143 | - | 2 | - | 16 | - | - |
| Hie mAb114 | - | 7 | - | 20 | - | - |
| Hie mAb114 UCA | - | 2 | - | - | - | - |
| Hie MEDI8852 | - | 2 | - | 15 | - | - |
| Hie MEDI8852 UCA | - | 6 | - | 20 | - | - |
| Hie REGN10987 | - | 8 | - | 13 | - | - |
| Hie S309 | - | 10 | - | 19 | - | - |
| Koenig G6 | 4275 | - | - | 4275 | - | - |
| Marks imm | - | - | 217 | - | - | - |
| Rosace Adalimumab | - | 14 | - | 14 | - | 14 |
| Rosace CD3022 | - | 6 | - | 6 | - | 6 |
| Rosace Golimumab | - | 5 | - | 5 | - | 5 |
| Shane. Trast. multi | - | - | - | 24 | - | - |
| Shane. Trast. zero | - | - | - | 422 | - | - |
| Warszawski D44 | - | - | - | 2049 | - | - |
| Wittrup CST | 274 | 137 | - | - | 822 | 411 |

Expression is measured in $\mu\text{g/mL}$ , thermostability with melting temperature, immunogenicity with percent of patients experiencing an anti-drug antibody response, binding with a dissociation constant $K_D$ , aggregation with wavelength shift, and polyreactivity with retention time.

### 3.2 Pipeline for Model Evaluation

We detail our pipeline for benchmarking protein language models in Supp. A.3. We used the antibody variable region sequence or structure as inputs for each model to assess their predictive capabilities, based off the corresponding model’s perplexity scores (averaged over all residues in the heavy and light chains). We report the Pearson (linear relationships, *r*), Spearman (monotonic relationships, *ρ*), and Kendall tau (ordinal relationships, *τ*) correlations to establish the connection between the model uncertainty values and the fitness metrics associated with the sequences in the dataset (Supp. A.5). If a protein language model correctly captures the biophysical landscapes of an antibody during training, it should assign higher confidence (low perplexity) to high fitness antibodies and low confidence (high perplexity) to low-fitness antibodies. All models were previously trained in their respective studies; we performed no additional fine-tuning prior to calculating perplexities.

Numerous computational models have been investigated for antibody design encompassing diverse approaches: (1) Decoder-only language models are trained using next-token prediction, and we investigate the ProGen2 suite, IgLM, and ProtGPT2; (2) encoder-only language models capture continuous representations of sequences, and we investigate AntiBERTy; and (3) inverse folding models predict protein sequences from structures, and we investigate ESM-IF and ProteinMPNN. To compare these deep learning methods versus physics-based models, we also calculated Rosetta energy for all sequences. Supp. A.4 provides an overview of all models tested and their corresponding (pseudo-)perplexity equations.

### 3.3 Fitness correlations with model perplexities

We asked whether model likelihoods, expressed as average perplexities, would correlate with experimentally measured fitness. In Fig. 1, we show two examples with the ProGen2-Small model. To summarize the correlations of all tested models over all datasets, we plotted the Pearson’s correlation coefficients (PCCs) in a heat map (Fig. 2; Spearman and Kendall tau coefficients are similar; see Supp. A.15). Supp. A.6 shows correlation plots for top performing models in each of the six fitness landscapes. ProGen2-Small obtained the most top performances (on seven datasets), with ProGen2-Medium, ProGen2-OAS, ESM-IF, and Rosetta Energy tied for second best (each are a top performer on six datasets). However, no model was a top performing model in all six fitness classes.

**Figure 1.**
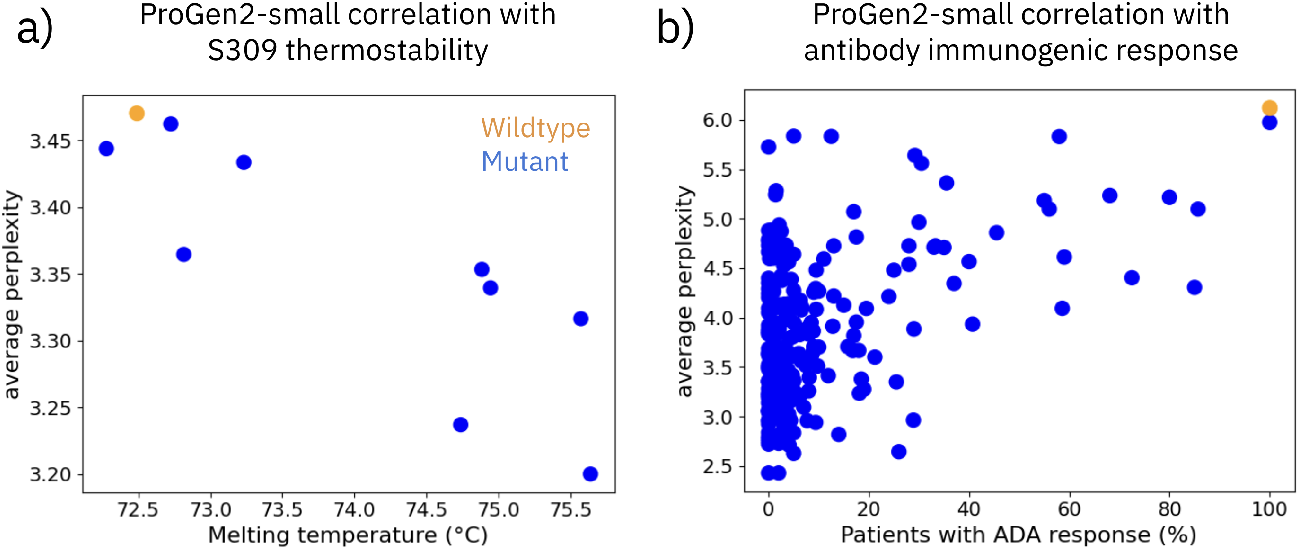
Examples of good and poor fitness prediction performance. (a) On a thermostability dataset of mutants of a patient-derived antibody that cross-neutralizes SARS-CoV-1 and 2, the language model correctly assigns higher confidence (lower perplexity) to the high melting temperature antibody variants (*r* = *−*0.84, *ρ* = −0.88, *τ* = −0.73). (b) On an immunogenicity dataset of percent anti-drug antibody responses (% ADA) from administered antibody therapeutics, the language model incorrectly assigns both high and low confidences to therapeutics that produce a 0% ADA response (*r* = 0.48, *ρ* = 0.32, *τ* = 0.23).

**Figure 2.**
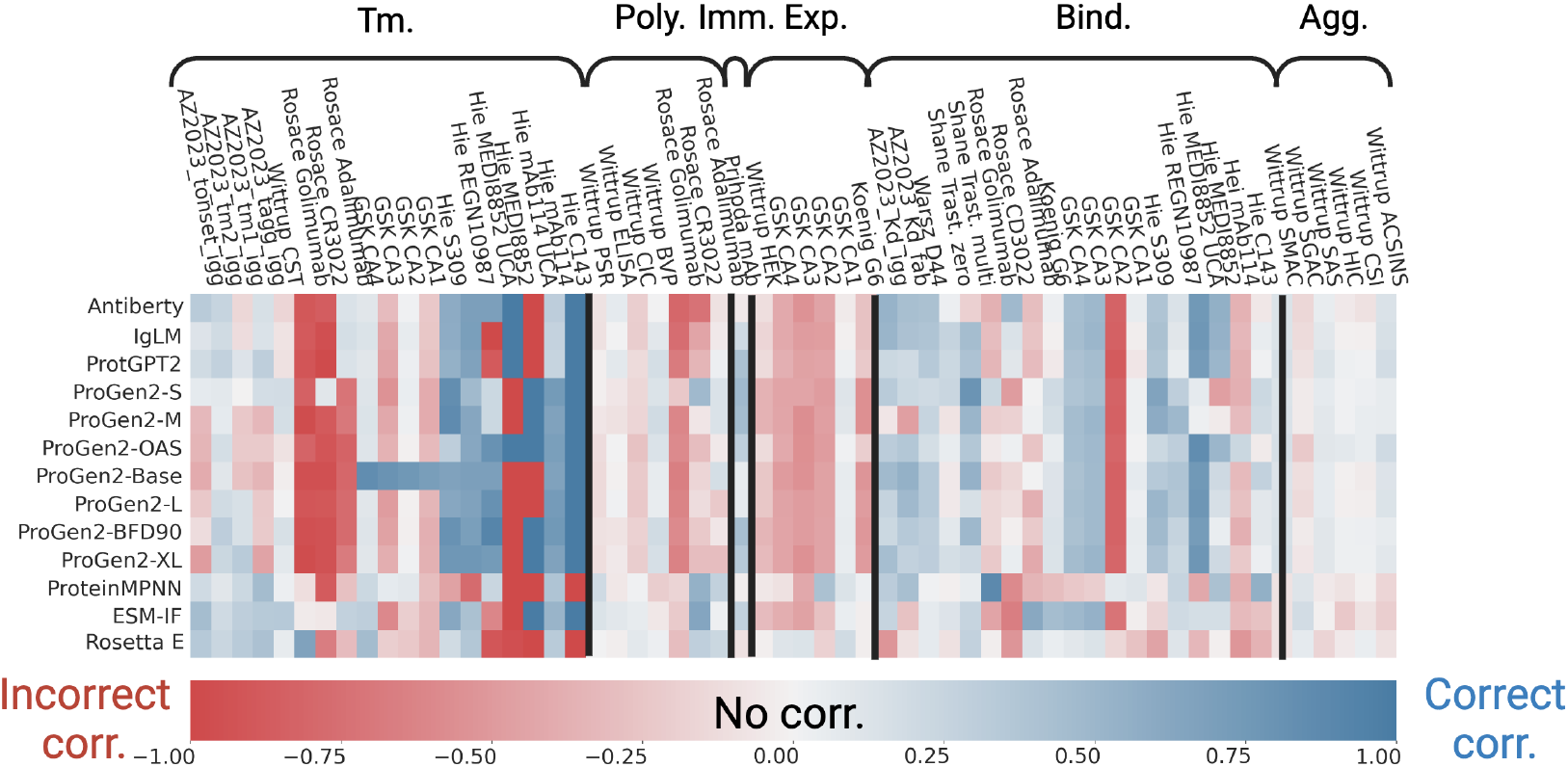
Summary of performance for each model-dataset pair. Pearson’s correlation coefficients (PCCs) for various protein model perplexities with a) aggregation, b) binding affinity, c) expression, d) immunogenicity, e) polyreactivity, and f) thermostability fitness prediction. The sign of the correlation was inverted for aggregation, expression, and thermostability, so that useful correlations will have positive PCC (blue).

### 3.4 Intrinsic biophysical properties are more accurately predicted than extrinsic properties

We next sought to identify different trends in the PCCs, such as whether models perform better on *intrinsic* properties, which are driven by inherent properties of the antibody (thermostability, aggregation), or *extrinsic* properties, which result from target biology and mechanism of action (expression, immunogenicity, binding affinity, polyreactivity). As shown in Supp. A.7, the absolute value of the PCC for all models was on average above 0.6, while it was significantly lower for binding affinity (< 0.4), expression (< 0.42), and immunogenicity (< 0.5). Thus, the intrinsic properties are better correlated with model likelihoods, which is unsurprising, since the models do not have access to contextual information.

### 3.5 Models are more accurate at distinguishing intra-family versus inter-family antibody sets

We next asked whether models were better at distinguishing multi-point mutants of antibodies originating from the same wild type (*intra-family*) or diverse antibodies from different wild type origins (*inter-family*). The absolute value of the PCC on the Hie *et al*. intra-family thermostability datasets is 0.77, while the thermostability prediction for the Jain *et al*. inter-family CSTs is 0.12 (Supp. A.8). The clinical stage therapeutics have each followed a different co-evolutionary maturity and selection processes, and therefore capturing these large sequence differences and properly assigning relatively nuanced fitness confidences may be more difficult than distinguishing sequences with less variability (e.g. single- and multi-point mutations). For the aggregation landscape we only have inter-family datasets (six from Jain *et al*.), and on average the average absolute PCC is below 0.2.

### 3.6 Parameter size impacts performance more that architecture and dataset composition

We also asked how correlations are affected by deep learning properties, e.g. architecture, dataset composition, and parameter size. We compared architectures by examining results of AntiBERTy (encoder-only language model) and IgLM (decoder-only language model), which are models trained on the same dataset of 558M antibody sequences from the Observed Antibody Space (OAS) [22]. For all six landscapes, AntiBERTy and IgLM display a similar performance, with the biggest variation being a greater range in correlations for polyreactivity datasets for AntiBERTy (Supp. A.9). A similar result was observed for dataset composition: When comparing three ProGen2 models with similar architecture yet distinct training datasets (ProGen2-OAS is trained on 554M antibody sequences, and ProGen2-Medium and -Base are trained on different compositions of UniRef90 and BFD30), no single model outperforms on all six landscapes (Supp. A.9). Prior studies reveal that an increase in model size typically leads to improved prediction performance [21, 18, 25]. In Supp. A.10, we plot the performance of four ProGen-2 models with increasing size: small (151M), medium (764M), large (2.7B), and xlarge (6.4B). While aggregation, binding affinity, expression, and immunogenicity prediction did not vary with model size, polyreactivity and thermostability improved noticeably. Thus, larger parameter sizes sometimes better captures the full complexity of the antibody fitness landscape.

### 3.7 Structure-based and sequence-based models perform similarly

We next asked whether explicitly providing structural information affect correlation performance. The sequence-based methods comprise AntiBERTy, IgLM, the ProGen2 suite, and the structure-based methods are ProteinMPNN, ESM-IF, and Rosetta Energy. Across all six fitness landscapes, sequence-based methods on average outperform the structure-based methods, with the most significant disparity being thermostability prediction (Supp. A.11). While sequence-based models must learn both structural syntactic and semantic mapping rules, structure-based methods already have the input encoded with structural interactions between CDRs and surrounding residues [5]. For the structure-based methods, no antigen information was provided, which could improve antibody fitness prediction in particular for the binding affinity landscape. Future work could predict the binding pose of each antibody mutant with their respective target antigen to score with structure-based methods.

### 3.8 Some models favor evolutionary signal rather than physical fitness

Finally, we investigated whether any models are biased towards evolutionary signal rather than physical protein fitness. The prevailing methods for protein structure prediction relies on an input protein representation coupled with a multiple sequence alignment (MSA) of homologous proteins to map evolutionary relationships between corresponding residues of genetically-related sequences. However, a language model that learns patterns in protein sequences across evolution may become biased towards evolutionary signal and assign higher fitness towards evolutionarily conserved mutations rather than evolutionarily divergent, possibly higher fitness mutations - a phenomenon that may be observed with some of the models benchmarked in FLAb. AntiBERTy, IgLM, and the entire ProGen2 suite assign higher confidence to the wild-type golimumab antibody, rather than the mutant antibody designs that have higher thermostability (Supp. A.12). However, physics-based Rosetta identifies the higher thermostability antibodies as more stable (lower Rosetta energy) than the wild type. Future work may consider encoding physics-based priors like Rosetta into a language model to negate evolutionary bias.

## 4 Conclusion

We constructed an antibody therapeutic property database and benchmarked the ability of widely adopted deep learning models to capture antibody properties. No model correlates well with all six properties, and model performance varies across datasets of the same property. While intrinsic biophysical properties are more readily captured, many struggle with extrinsic properties like expression, immunogenicity, binding affinity, and polyreactivity. Additionally, the number of learnable parameters seems to influence performance more than the model pretraining data composition or architecture. Promising directions for protein language models involve incorporating protein structure, antigen information, physics-based priors, or the ever-growing antibody fitness data in the model.

Unfortunately, there are still too few data points in these datasets for training new models (a recent study estimates that at least 10^4^ binding affinities are needed for the binding affinity prediction task [14]). In practice, we will need more nuanced metrics than any single model’s likelihoods, since they should not be expected to capture all diverse fitness metrics from immunogenicity to binding affinity. Looking toward the discovery of new data and the development of new models, we invite contributions to FLAb toward working to the goal of achieving reliable, well-behaved antibody therapeutics from computational designs.

## A Supplementary material

### A.1 Dataset and code availablility

Data and model scoring methods used for benchmarking in FLAb can be accessed at https://github.com/Graylab/FLAb.

### A.2 Landscapes and dataset descriptions

**Figure 1.**
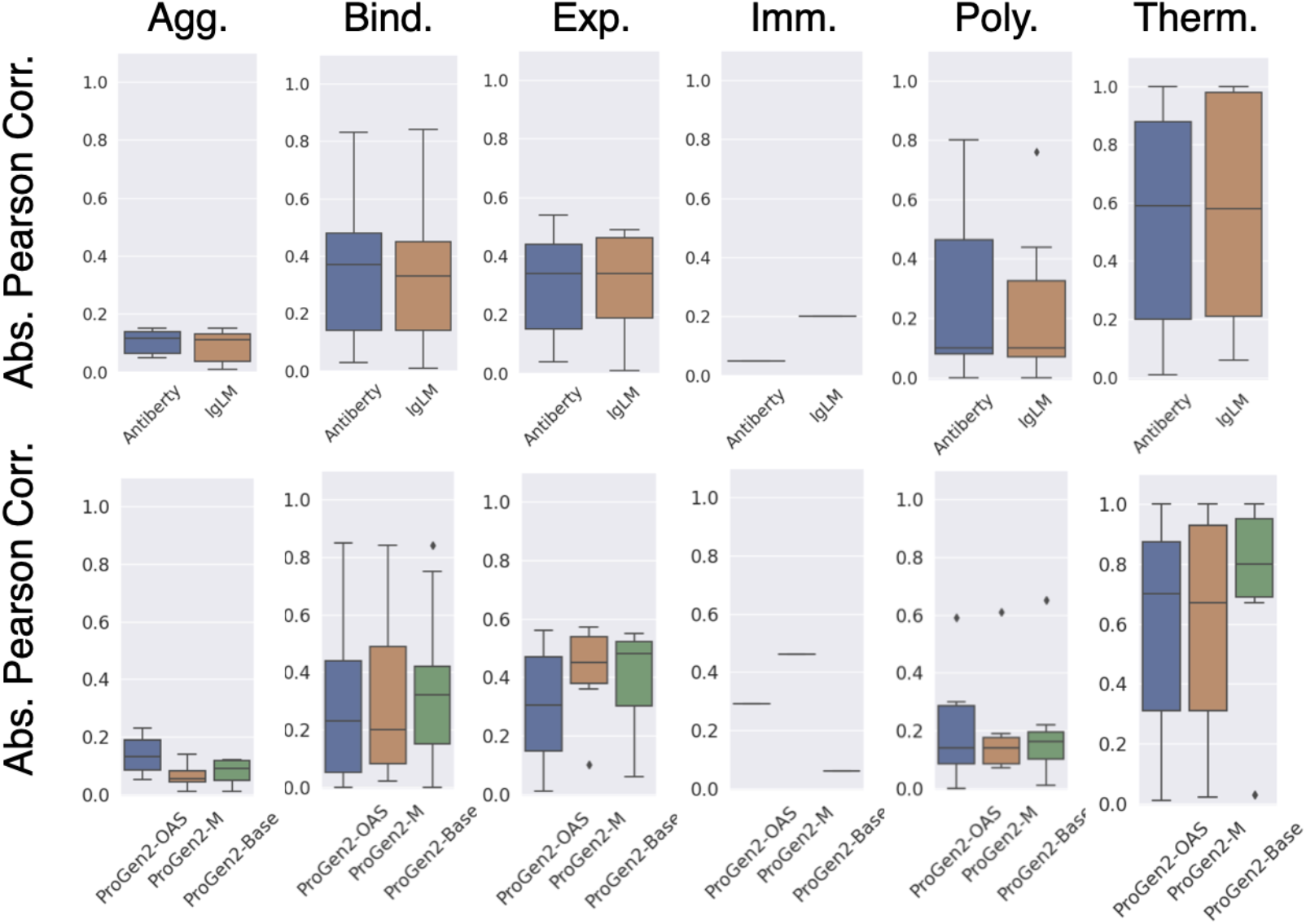
Six classes of biophysical data relevant to antibody developability. Desirable values for each property is shown in green, while undesirable values are shown in red. (a) Antibody expression is assessed using ELISA fluorescent signal, with high expression indicating high optical density (1.0-1.5). (b) Thermostability is assessed through differential scanning calorimetry, which measures the heat capacity change of a sample with temperature, where temperatures above 70°C are considered ideal. (c) Immunogenicity was quantified as the percentage of patients developing anti-drug antibodies (ADA) in response to therapeutic administration, where an ideal, non-immunogenic antibody results in 0% ADA response. (d) Binding affinity is assessed using the equilibrium dissociation constant (*K*_*d*_), with a desirable *K*_*d*_ typically falling in the low nanomolar to picomolar range. (e) Aggregation can be measured using an AC-SINs assay, where no change change in the measured plasmon wavelength shift is ideal. (e) Polyreactivity can be measured with CIC retention time, where therapeutic antibodies are expected to have a retention time of at least 10 minutes.

*Expression* ensures the production of antibodies in a host cell system, which is necessary to isolate a molecule for further testing and directly affects production yield and cost of manufacturing. The enrichment ratio quantifies the expression of each variant antibody compared to a wildtype antibody. The largest set of expression data is from Koenig *et al*., who conducted an extensive mutational analysis over 4275 mutations at all positions within the variable domain of a high-affinity anti-VEGF antibody (G6.31) [7]. We also analyze 4 sets of designed antibodies (CA1, CA2, CA3, and CA4) from GlaxoSmithKline and our lab, and expression titer in HEK cells for a list of clinical stage therapeutic (CST) antibodies [6].

*Thermostability* ensures an antibody will maintain its structure and function when exposed to heat, particularly during manufacturing, storage, and administration. Antibodies with high thermostability are more likely to remain potent over extended periods and under different storage conditions. A diverse set of thermostability measures come from Hie *et al*. who employed language model-guided evolution techniques to investigate mutations in seven antibodies. This set comprised four clinically relevant and highly matured antibodies (MEDI8852, mAb114, S309, and REGN10987), as well as three unmatured antibodies (MEDI8852 UCA, mAb114 UCA, and C143), providing a set of melting temperature values for mutants of the set of evolved antibodies [4]. We also provide thermostability data for the aformentioned GSK antibodies and Adalimumab, CD3022, Golimumab from Rosace *et al*.[10].

*Immunogenicity* refers to the ability of a therapeutic antibody to elicit an undesirable immune response in the body, leading to the generation of anti-drug antibodies (ADAs). ADAs can recognize and neutralize therapeutic antibodies, reducing their efficacy and potentially causing adverse effects. Minimizing immunogenicity is important for therapeutic antibodies to maintain their efficacy and safety. Marks *et al*. provides a dataset of 198 human, 229 humanized, 63 chimeric, and 13 murine antibody sequences, as well as reported anti-drug antibody (ADA) responses from patients for 217 therapeutics [8].

*Binding affinity* ensures the prolonged physical contact during an interaction between an antibody and target antigen, impacting their ability to block pathways or target disease molecules. GSK, Hie *et al*., and Rosace *et al*. provide a combined 13 sets of antibody binding affinity data. Warszawski *et al*. aimed to investigate the mutational tolerance of 135 positions within the anti-lysozyme antibody D44.1, for a total of 2048 mutants [15], and the Koenig *et al*. G6.31 mutant dataset provides 4275 data points for binding [7]. Shanehsazzadeh *et al*. redesign the trastuzumab antibody with 442 zero-shot mutants and 24 multi-step mutants [13].

*Polyreactivity* of an antibody allows it to bind to multiple antigens. In the context of therapeutic antibodies, although polyreactivity can sometimes be beneficial (if it is desired to bind to multiple targets) or problematic (if the antibody interfers with normal cellualar function due to off-target binding). Rosace *et al*. provide polyreactivity data for the Adalimumab, CD3022, and Golimumab variants [10], and Wittrup *et al*. provide polyreactivity measurements using BVP, CIC, ELISA, and PSR assays on CSTs [6].

*Aggregation* refers to the process of individual antibodies coming together to form larger assemblies, or aggregates. Aggregation can be problematic as it leads to reduced therapeutic efficacy and potentially harmful immune responses. The only aggregation data is 822 fitness values from AC-SINS, CSI, HIC, SAS, SGAC, and SMAC assays on CSTs [6].

### A.3 Scoring pipeline

**Figure 2.**
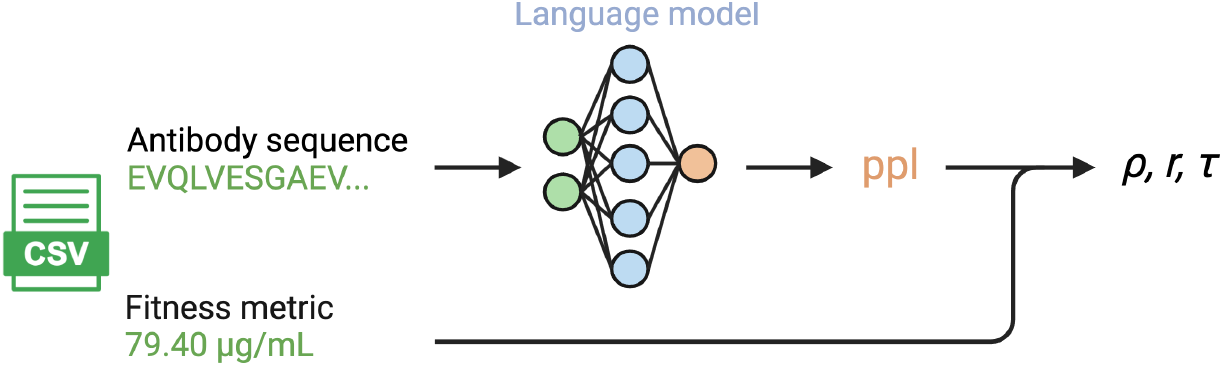
Pipeline for benchmarking protein language models. All fitness datasets contain columns for antibody heavy chain sequence, antibody light chain sequence, and an associated fitness metric. For each protein language model, we separately input the heavy and light sequence to return two perplexity scores, and we tabulate the average perplexity between the two sequences. For structure-conditioned language models, we first predict the antibody structure with IgFold [11], and then tabulate the single perplexity scored from the model. Correlation metrics (Pearson’s, Spearman’s and Kendall tau’s correlations) are calculated between average perplexity and the fitness measure. No antigen information is provided for any benchmarked models.

### A.4 Classes of language models scored

#### A.4.1 Decoder-only language models

Decoder-only language models have proven to be effective in generating plausible and novel protein sequences. These models are trained using a next-token prediction objective, where the probability of the next amino acid is influenced by the entire preceding sequence. During training, a database of sequences is utilized to predict *P* (s_*i*_|s_*<i*_), enhancing the model’s ability to generate accurate sequences.

We evaluate the zero-shot prediction of therapeutic properties by correlating to the perplexity of each sequence under those models:

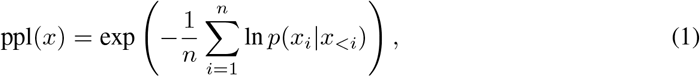

where *x* = (*x*_1_, *x*_2_, …, *x*_*n*_) is a sequence consisting of *n* tokens.

The decoder models we benchmark are **ProGen2** [9], **IgLM** [14], and **ProtGPT2** [3]. The ProGen2 models come in various sizes, ranging from 151M to 6.4B parameters, pretrained on a mixture of UniRef90 and BFD90 databases. IgLM formulates the design task based on text-infilling using a standard left-to-right decoder (GPT-2), trained on a non-redundant set of 558M antibody sequences obtained from the Open Antibody Sequence (OAS) database. **ProtGPT2** is a 738M parameter model trained on 50M non-annotatedsequences spanning the entire protein space [3].

#### A.4.2 Encoder-only language models

Encoder-only language models capture comprehensive information in a continuous abstract representation. These models are utilized for this purpose, allowing the learning of representations that can be broadly applied. A subset of residues is randomly chosen and replaced with a special mask token. The model is then trained to predict the identities of these masked residues.

In an encoder-only model, an estimation of perplexity can be obtained by calculating the exponential of the negative pseudo-log-likelihood, or pseudo-perplexity:

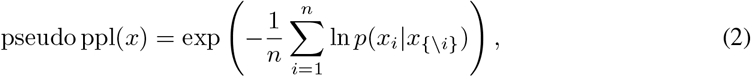

where *x*_*{-i}*_ is the set of all residues except *x*_*i*_.

In this category of models we focus on **AntiBERTy**, a 26M parameter model pretrained on 558M natural antibody sequences from OAS [12].

#### A.4.3 Structure-conditioned language and network models

Generative deep learning architectures that predict protein sequences from structures are known as structure-conditioned language models. Like decoder-only language models, for structure-encoded models we can evaluate the perplexity for each antibody sequence-structure pair:

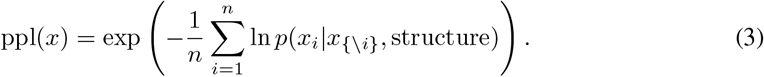

**ESM-IF** uses an autoregressive encoder-decoder architecture, where the model is tasked with recovering the native sequence of protein from the coordinates of its backbone atoms [5]. **ProteinMPNN** uses a message-passing neural network with 1.4M parameters that predicts protein sequences using several protein backbone geometry [2]. Structures of all antibody mutants are predicted with IgFold [11] prior to scoring with inverse folding models.

#### A.4.4 Physics-based models

We seek to compare the performance of protein language models mentioned in 4.1 - 4.3 versus empirical models of protein energy, which has been a longstanding approach for protein design efforts. **Rosetta**, the classic protein structure prediction and design software, employs an optimized energy function ref2015 that assesses the energy of atomic interactions within a globular protein [1]. Score functions within Rosetta are composed of weighted sums of various energy terms. Some of these terms correspond to physical forces, such as electrostatics and Van der Waals (VdW) interactions, while others represent statistical terms, like the likelihood of observing specific torsion angles in Ramachandran space:

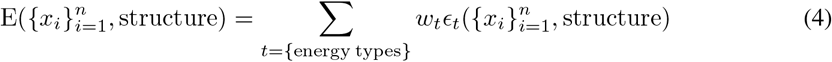

Where *ϵ*_*i*_ is a Rosetta energy term, and *w*_*i*_ is the respective weighted number. Rosetta’s energy calculation does not directly correspond to physical energy units, and are instead expressed in Rosetta energy units. A lower score indicates a higher likelihood of the structure being closer to the native structure. Structures of all antibody mutants are predicted with IgFold prior to calculating Rosetta energy.

### A.5 Statistical correlation

After obtaining predicted scores from each of the benchmarked models, we used a set of one linear and two non-linear correlation metrics to determine what relationships exist with the respective fitness dataset. Pearson’s correlation coefficient measures the strength and direction of the linear relationship between two variables, defined as:

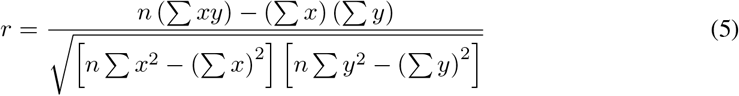

where *rϵ*[*−*1, +1], *n* is the number of data points, *x* is the fitness measurement and *y* is perplexity. We also calculate Spearman’s correlation coefficient, which captures the strength and direction of the monotonic relationship between two variables, defined as:

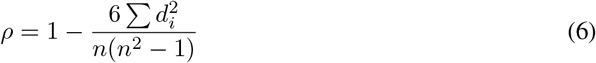

where ρ*ϵ*[*−*1, +1], *n* is the number of data points, and 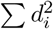 is the squared difference between the ranks of variables *x* and *y*.

Kendall’s tau coefficient is used to quantify the strength and direction of the ordinal relationship between two variables, defined as:

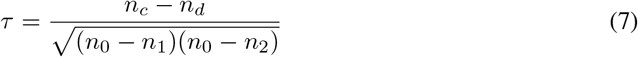

Where *τϵ*[*−*1, +1], *n*_0_ is the total number of pairs of data points, *n*_1_ is the number of pairs that have tied values for the first variable, *n*_2_ is the number of pairs that have tied values for the second variable, *n*_*c*_ is the number of concordant pairs of data points, and *n*_*d*_ is the number of discordant pairs of data points.

### A.6 Overview of top performing models

**Figure 3.**
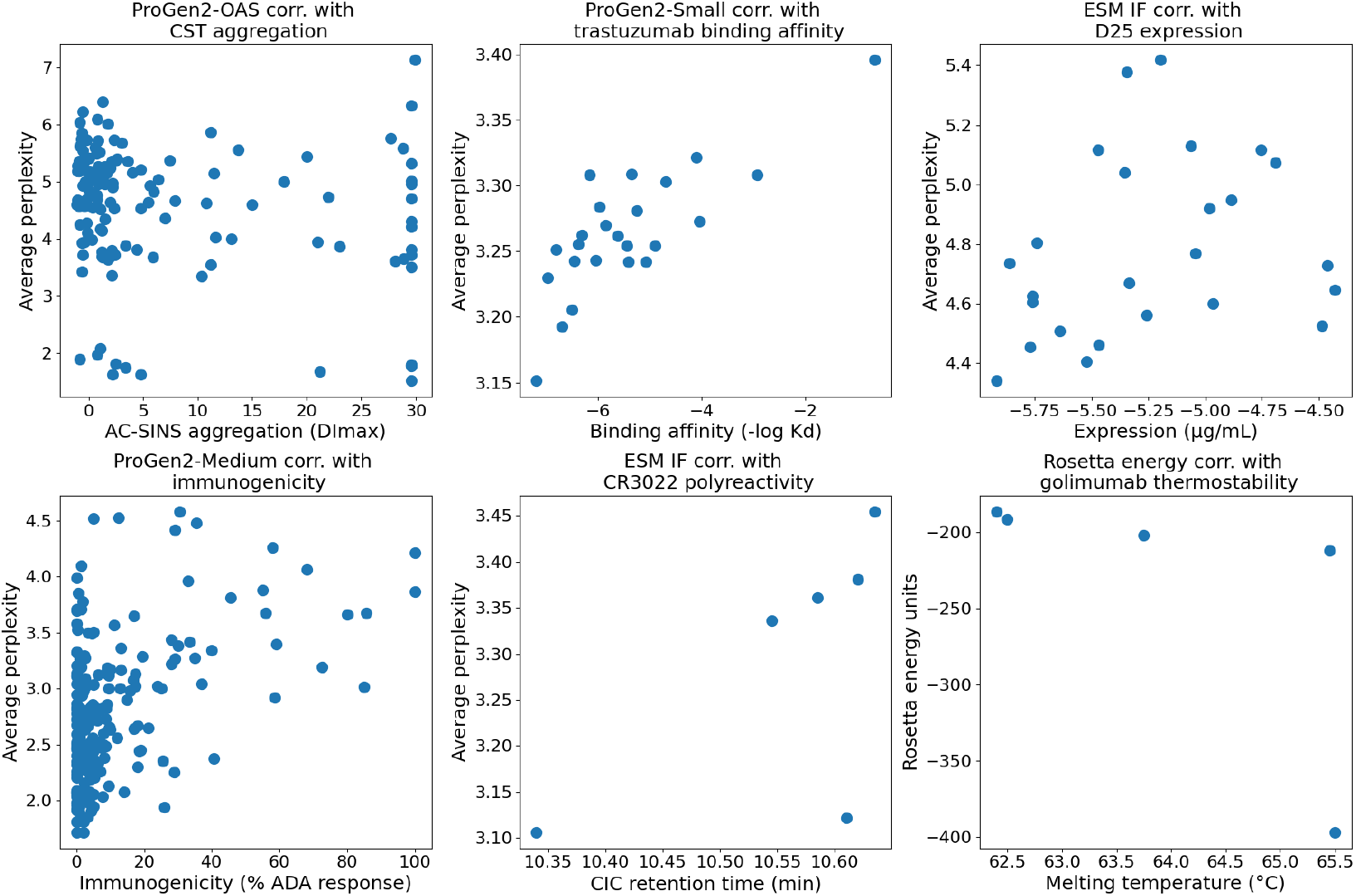
Performance plots for top performing model in each fitness landscape. From left to right across each row, the top performing model for the CST AC-SINS aggregation dataset was ProGen2-OAS; for the trastuzumab binding affinity dataset was ProGen2-Small; for the D25 expression dataset was ESM-IF; for the mAb immunogenicity dataset was ProGen2-Medium; for the CD3022 polyreactivity dataset was ESM-IF; and for the golimumab thermostability dataset was Rosetta energy.

### A.7 Intrinsic biophysical properties are more accurately predicted than extrinsic

**Figure 4.**
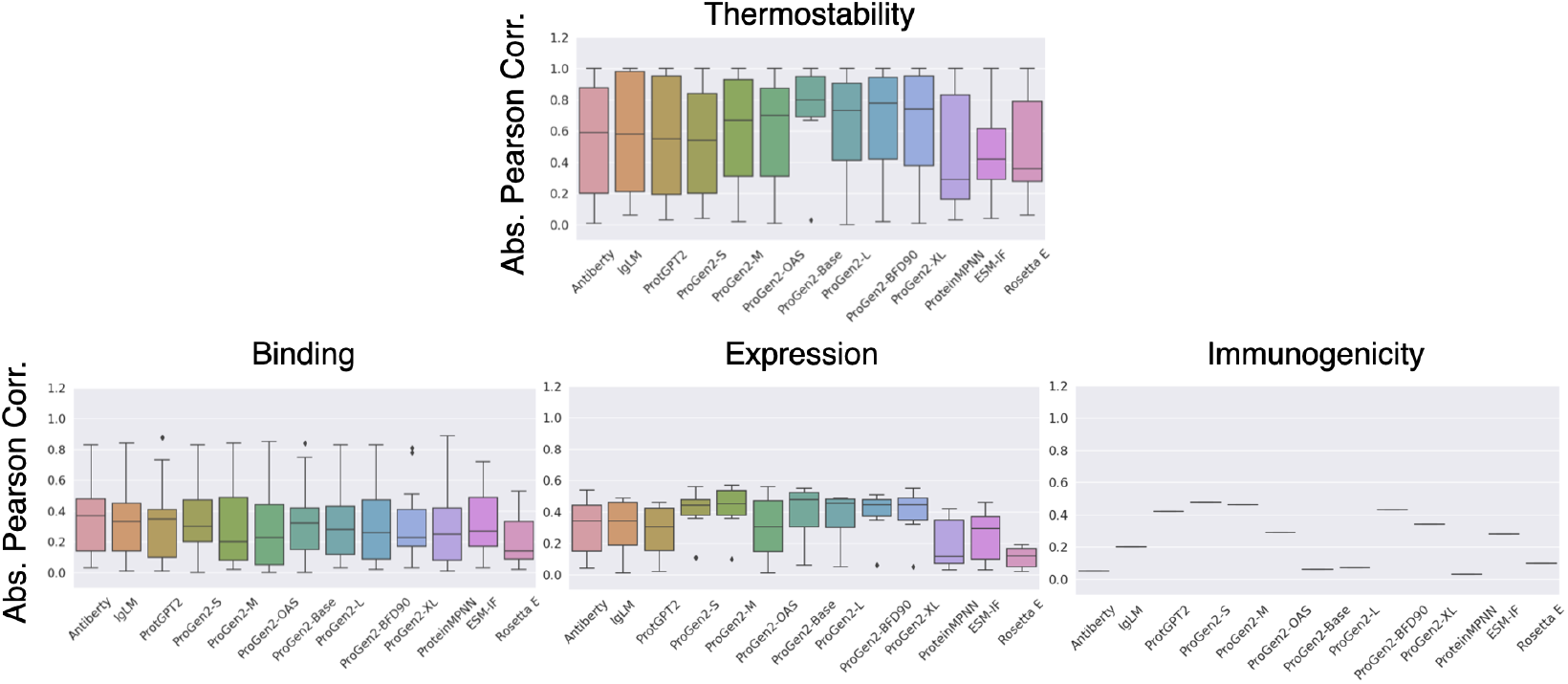
Comparison of performance on intrinsic and extrinsic biophysical property prediction. Intrinsic properties are impacted by inherent properties of the antibody, while extrinsic properties result from target biology and mechanisms of action. It can be seen that models better predict fitness variations in intrinsic properties.

### A.8 Models are more accurate at distinguishing intra-family versus inter-family antibody datasets

**Figure 5.**
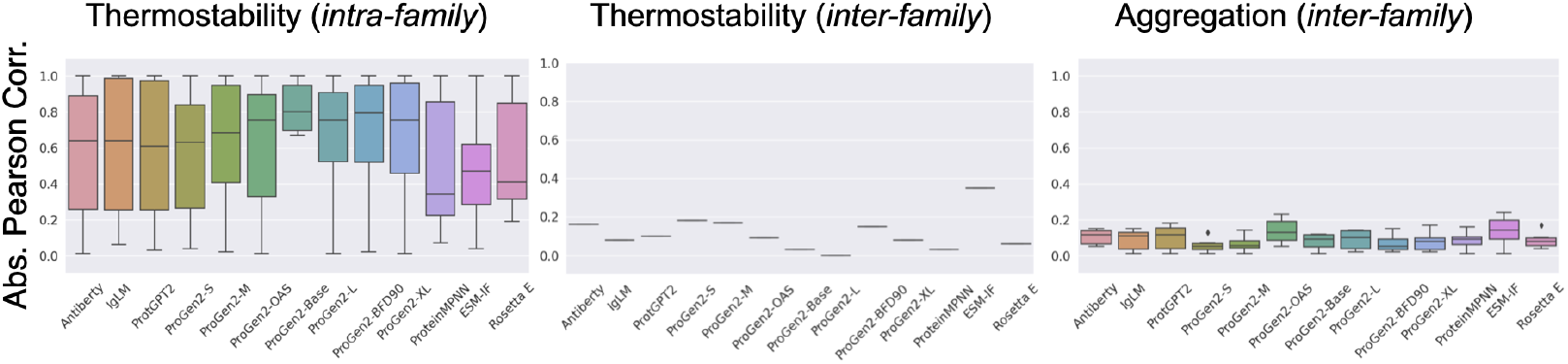
Comparison of performance on intra-family and inter-family datasets. Models are more accurate at distinguishing intra-family versus inter-family antibody sets. It can be seen that models better predict fitness variations in intra-family datasets.

### A.9 Architecture and dataset composition deviations do not significantly impact performance

**Figure 6.**
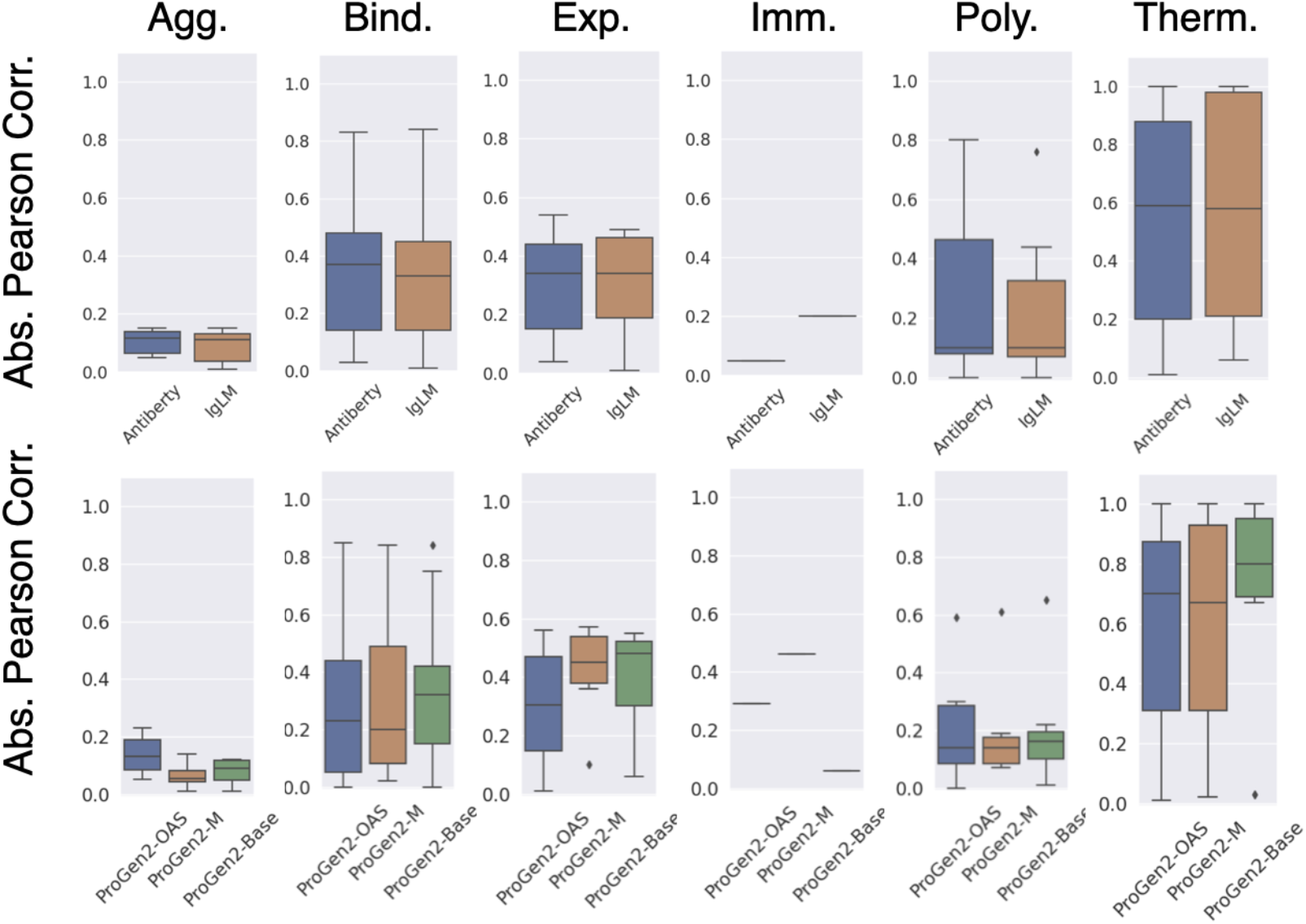
Comparison of performance on architecture and data composition variations. AntiB-ERTy and IgLM are trained on the same dataset of 558M antibodies, allowing for a comparison of architectural differences. ProGen2-OAS, Medium, and Base are 764M parameter models trained on different datasets, allowing for a direct comparison of dataset differences. In both instances, architectural and dataset differences do not seem to significantly impact performance.

### A.10 Parameter size influences performance over architecture and dataset composition

**Figure 7.**
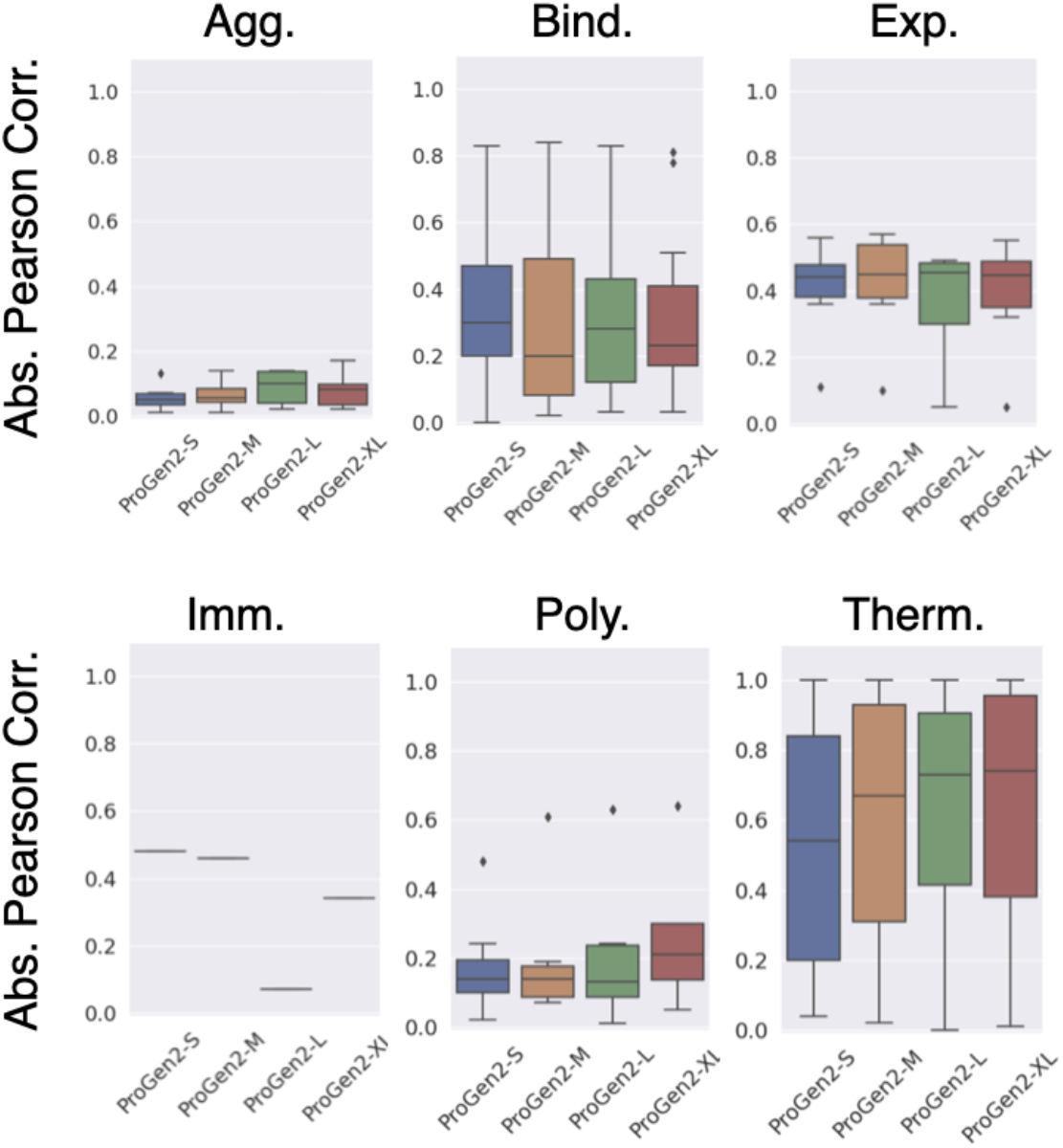
Comparison of performance on parameter size variations. ProGen2 introduced several models with varying parameter sizes, including Small (151M), Medium (764M), Large (2.7B), and XLarge (6.4B) - allowing for a comparison of the effect of parameter size on performance.

### A.11 Structure-based and sequence-based models perform similarly

**Figure 8.**
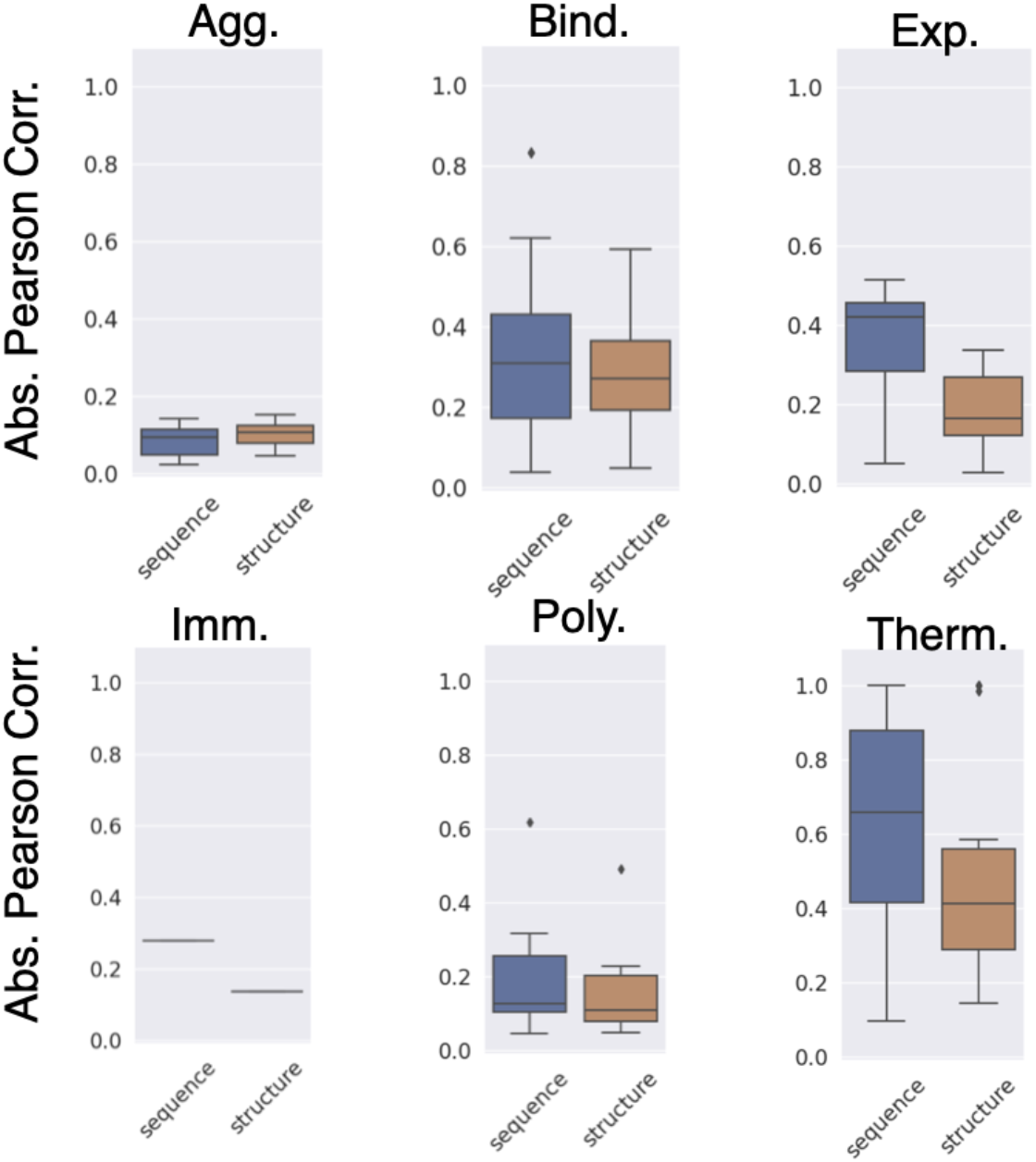
Comparison of performance on structure-based and sequence-based models. Sequence-based models bar plots indicate the average performance across AntiBERTy, IgLM, ProtGPT2, and ProGen2. Structure-based models bar plots indicate the average performance across ProteinMPNN, ESM-IF, and Rosetta.

### A.12 Some models favor evolutionary signal rather than physical fitness

**Figure 9.**
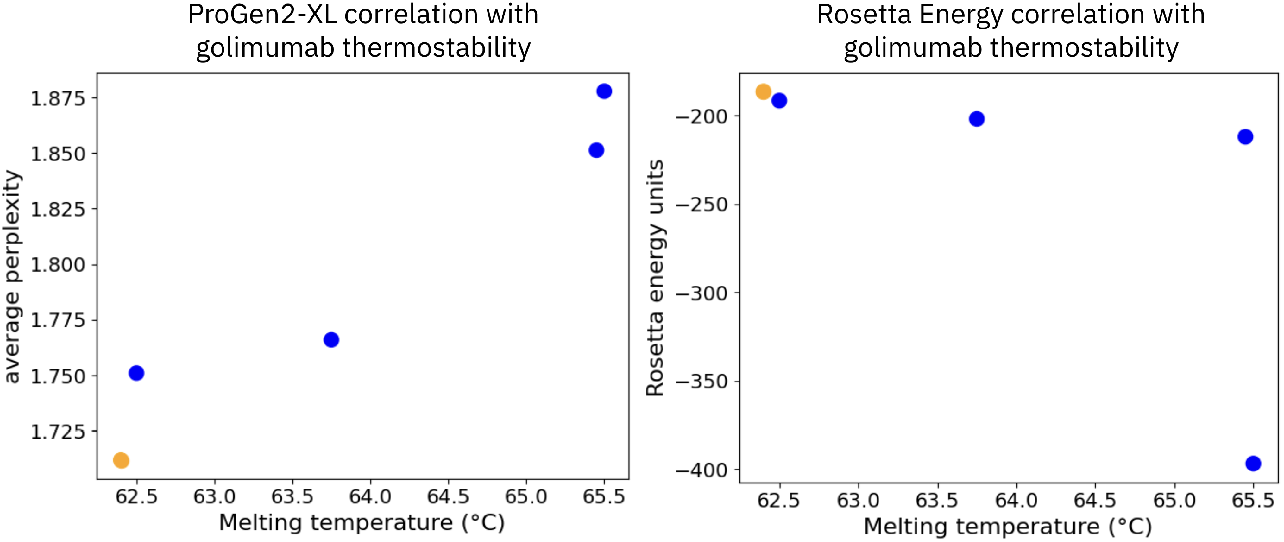
Protein language models might exhibit bias towards evolutionary signal. Displayed are the scores of ProGen2-XL and Rosetta energy for mutants of clinically approved Golimumab. It can be seen that the language model incorrectly assigns higher confidence to the wildtype antibody, while physics-based Rosetta correctly assigns higher stability to the more thermostable Golimumab variants.

### A.13 Glossary of terms

**Antibody**. Antibodies, also known as immunoglobulins, are Y-shaped proteins produced by specialized white blood cells called B cells. They play a crucial role in the immune system by recognizing and binding to specific foreign substances, called antigens, such as pathogens or toxins. This binding marks the antigens for destruction by other immune cells. The primary region of variability and binding, the Fv region, consists of a heavy chain and light chain sequence.

**CDRs**. Complementarity-determining regions (CDRs) are short stretches of amino acids within the variable regions of antibodies. These loops are responsible for directly interacting with antigens. By altering their conformation, CDR loops create a unique antigen-binding site, allowing antibodies to recognize a diverse array of antigens.

**Therapeutic antibody**. Therapeutic antibodies are antibodies that are designed or engineered for medical use. They can be utilized to treat various diseases, including cancers, autoimmune disorders, and infectious diseases, by targeting specific molecules involved in these conditions.

**Developability**. Antibody developability refers to the set of biological biophysical characteristics that determines it’s potential to be manufactured and perform it’s therapeutic objective in a patient. These characteristics include high-level expression, high solubility, covalent integrity, conformational and colloidal stability, low polyspecificity, and low immunogenicity.

**Protein fitness**. Protein fitness refers to the ability of a protein to perform its intended biological functions effectively. A protein’s fitness is multi-dimensional and context-dependent, determined by its structure, stability, and interactions with other molecules. Proteins with higher fitness are more likely to contribute positively to cellular processes.

**Fitness landscape**. The fitness landscape of proteins represents the relationship between protein variations (mutations) and their corresponding fitness levels. It describes how different mutations can impact a protein’s function, stability, and interactions within a biological context.

**Thermostability**. Thermostability refers to an antibody’s ability to maintain its structure and function when exposed to elevated temperatures. Antibodies with high thermostability are more resilient and can have longer shelf lives.

**Expression**. Antibody expression is the process by which cells, often genetically engineered, produce antibodies. This can occur within organisms or in laboratory settings. Efficient expression is crucial for generating sufficient quantities of antibodies for research or therapeutic purposes.

**Immunogenicity**. Immunogenicity refers to the likelihood of an antibody itself inducing an immune response when introduced into an organism. Overly immunogenic antibodies might trigger adverse reactions in patients.

**Binding affinity**. The binding affinity of antibodies defines how strongly an antibody interacts with its target antigen. A high binding affinity implies a strong and specific interaction, which is desirable for effective antigen recognition and neutralization.

**Polyreactivity**. Polyreactivity is the ability of an antibody to bind to a variety of self and foreign anti-gens, which may be completely unrelated, and is often attributed to a more conformationally flexible antigen binding pocket.

**Aggregation**. Aggregation of antibodies refers to the process by which individual antibody molecules come together to form large complexes or aggregates. These aggregates can reduce efficacy, trigger an immune response, or affect the storage stability after manufacturing.

### A.14 Limitations

A limitation to this work is that the available labeled antibody fitness datasets are currently small. Many of these datasets consist of a relatively small number of data points, often containing fewer than 30 data points. While these datasets provide insights into the prediction capabilities of AI models, the limited data points present challenges in establishing robust correlations between true and predicted fitness. Out of 1872 of the calculated correlations (52 datasets, 12 models, and 3 correlations per dataset-model pair), only 515 correlations had an associated p-value less than 0.05. Nevertheless, this benchmark provides a starting point for assessing the predictive potential of AI models in the realm of therapeutic antibody fitness. The results obtained offer crucial insights into the strengths and weaknesses of different approaches, guiding future research efforts towards enhancing predictive accuracy and robustness. Additionally, we call upon antibody engineers in academia and industry to generate additional data and contribute it to this repository. Future work correlating antibody embeddings with these properties must use caution with the currently small size of FLAb.

Additionally, since the fitness datasets we provide contain experimental data from independent studies, the exact conditions for each experiment are likely different from one study to another, ultimately leading to inconsistencies in the reported experimental data (e.g., binding affinities may be affected by different solution concentrations in each study). The limited availability of public experimental data on therapeutic antibody candidates hinders the comprehensive evaluation of protein language models and development of novel protein design models. We urge collaboration between experimentalists and computational scientists to share therapeutic antibody data, enabling thorough analysis and improving the therapeutic antibody design process.

### A.15 Summarizing heat map of statistical correlations

**Figure 10.**
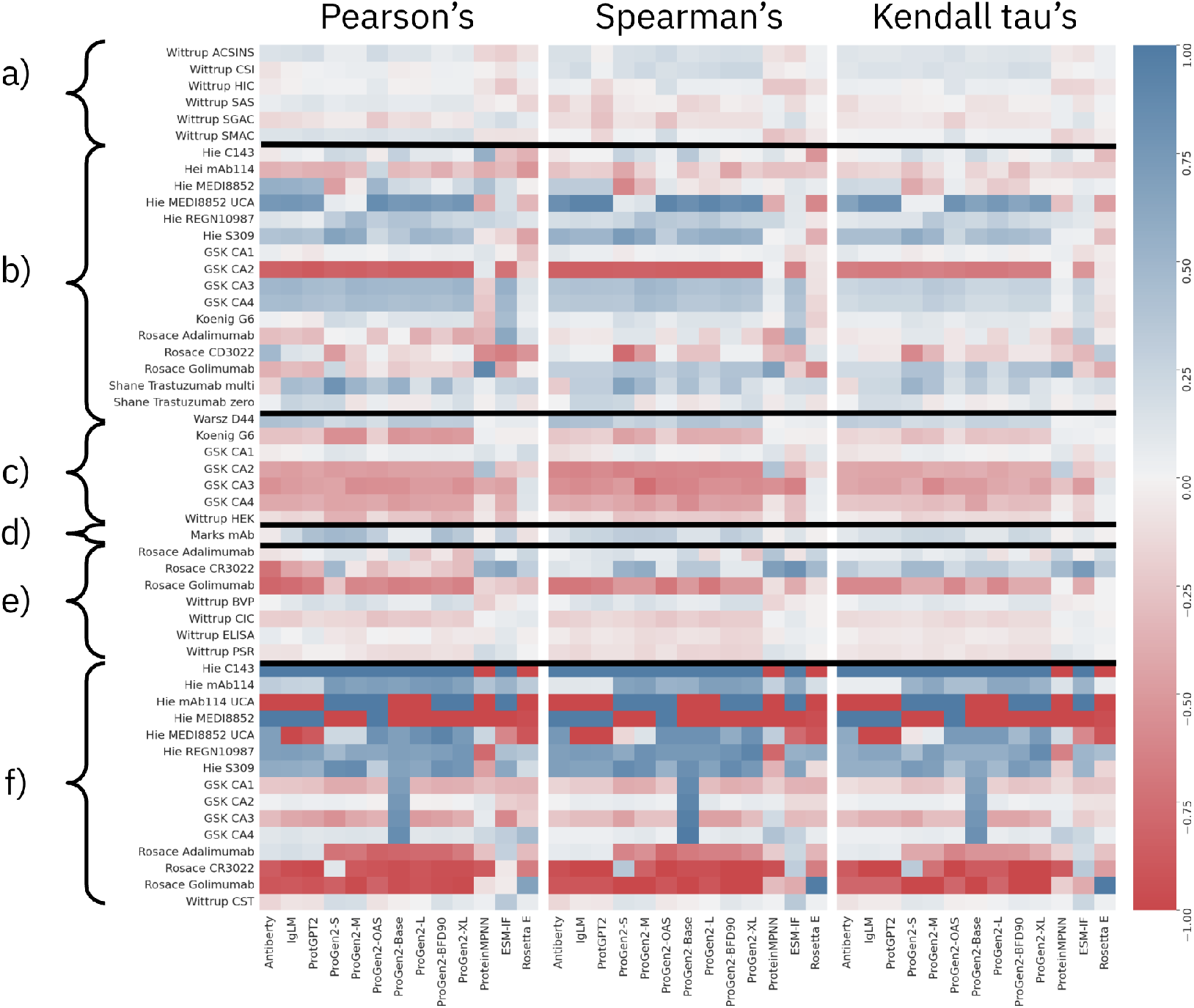
Summary of performances for each model-dataset pair. Linear (Pearsons’s) and non-linear (Spearman’s, Kendall tau’s) correlations are provided for a) aggregation, b) binding affinity, c) expression, d) immunogenicity, e) polyreactivity, and f) thermostability fitness prediction. Models generally perform best with thermostability and binding affinity datasets of single point mutants, but struggle with aggregation and expression datasets of antibodies with differing wildtype origins.

**Table 1.** Summary of Pearson correlations. Correlations with statistical significance (p < 0.05) are shown in bold.

| Dataset | AntiBERTy | IgLM | ProTGPT2 | Small | Medium | OAS | ProGen2<br>Base | Large | BFD90 | Xlarge | proteinMPNN | ESM IF | Rosetta |
| --- | --- | --- | --- | --- | --- | --- | --- | --- | --- | --- | --- | --- | --- |
| Wittrup ACSINS Agg. | -0.14 | -0.15 | -0.18 | -0.06 | -0.06 | -0.20 | -0.12 | -0.13 | -0.06 | -0.09 | 0.16 | **0.20** | 0.10 |
| Wittrup CSI Agg. | 0.10 | 0.02 | -0.01 | -0.03 | -0.05 | -0.05 | 0.04 | 0.03 | -0.03 | -0.02 | 0.01 | 0.09 | -0.04 |
| Wittrup HIC Agg. | 0.05 | 0.01 | -0.02 | -0.01 | -0.01 | -0.08 | 0.01 | 0.02 | -0.02 | -0.02 | 0.09 | **0.24** | 0.05 |
| Wittrup SAS Agg. | -0.05 | -0.09 | -0.13 | -0.07 | -0.09 | -0.10 | -0.07 | -0.07 | -0.10 | -0.10 | 0.05 | 0.01 | **0.17** |
| Wittrup SGAC Agg. | 0.15 | 0.13 | 0.10 | 0.04 | 0.04 | 0.23 | 0.11 | 0.14 | 0.04 | 0.07 | -0.11 | **-0.18** | -0.07 |
| Wittrup SMAC Agg. | -0.13 | -0.13 | -0.16 | -0.13 | -0.14 | -0.16 | -0.12 | -0.14 | -0.15 | -0.17 | 0.09 | 0.10 | 0.09 |
| Hie C143 Kd | -0.06 | 0.05 | 0.01 | 0.22 | 0.19 | -0.04 | 0.28 | 0.09 | 0.09 | 0.03 | **0.56** | -0.27 | -0.36 |
| Hie mAb114 Kd | -0.36 | -0.33 | -0.37 | -0.38 | -0.31 | 0.23 | -0.32 | -0.28 | -0.38 | -0.23 | -0.39 | -0.32 | **-0.52** |
| Hie MED18852 Kd | **0.56** | **0.54** | 0.50 | -0.47 | -0.04 | 0.50 | 0.21 | 0.16 | 0.04 | 0.21 | 0.42 | 0.17 | -0.02 |
| Hie MED18852 UCA Kd | **0.75** | **0.77** | 0.73 | 0.07 | 0.02 | 0.85 | **0.75** | **0.78** | **0.72** | **0.78** | -0.42 | 0.17 | -0.33 |
| Hie REGN10987 Kd | 0.14 | 0.10 | 0.05 | 0.30 | 0.49 | 0.30 | 0.39 | 0.37 | 0.26 | 0.22 | 0.25 | 0.20 | 0.11 |
| Hie S309 Kd | 0.39 | 0.42 | 0.38 | **0.66** | **0.59** | 0.22 | 0.35 | 0.50 | **0.47** | 0.40 | -0.07 | -0.17 | -0.45 |
| GSK CA1 Kd | 0.05 | -0.03 | -0.07 | 0.02 | 0.02 | 0.00 | 0.07 | 0.03 | 0.06 | 0.05 | 0.11 | -0.03 | -0.27 |
| GSK CA2 Kd | **-0.83** | **-0.84** | **-0.88** | **-0.83** | **-0.84** | -0.81 | **-0.84** | **-0.83** | **-0.83** | **-0.81** | 0.08 | **-0.72** | -0.15 |
| GSK CS3 Kd | **0.49** | 0.50 | 0.45 | 0.43 | 0.50 | 0.48 | 0.47 | 0.47 | 0.47 | 0.51 | -0.20 | 0.54 | 0.13 |
| GSK CA4 Kd | 0.44 | 0.45 | 0.41 | 0.39 | 0.45 | 0.44 | 0.42 | 0.43 | 0.42 | 0.46 | -0.24 | 0.49 | 0.09 |
| Koenig G6 Kd | **0.03** | -0.01 | **-0.05** | **0.20** | **0.17** | 0.05 | **0.12** | **0.12** | **0.14** | **0.13** | -0.29 | 0.45 | 0.04 |
| Rosace Adalimumab Kd | -0.29 | -0.26 | -0.31 | 0.00 | 0.02 | -0.26 | 0.00 | -0.38 | -0.22 | -0.41 | -0.35 | **0.61** | 0.14 |
| Rosace CD3022 Kd | 0.48 | 0.14 | 0.10 | -0.49 | -0.18 | 0.02 | -0.15 | -0.11 | -0.02 | 0.04 | -0.59 | -0.66 | -0.53 |
| Rosace Golinumab Kd | -0.37 | -0.30 | -0.34 | 0.23 | -0.20 | -0.05 | -0.22 | -0.15 | -0.15 | -0.23 | **0.89** | -0.44 | -0.03 |
| Warsz D44 Kd | **0.42** | **0.39** | 0.35 | **0.20** | **0.25** | 0.17 | **0.35** | **0.35** | **0.30** | 0.31 | 0.01 | 0.07 | 0.07 |

**Table 2.**
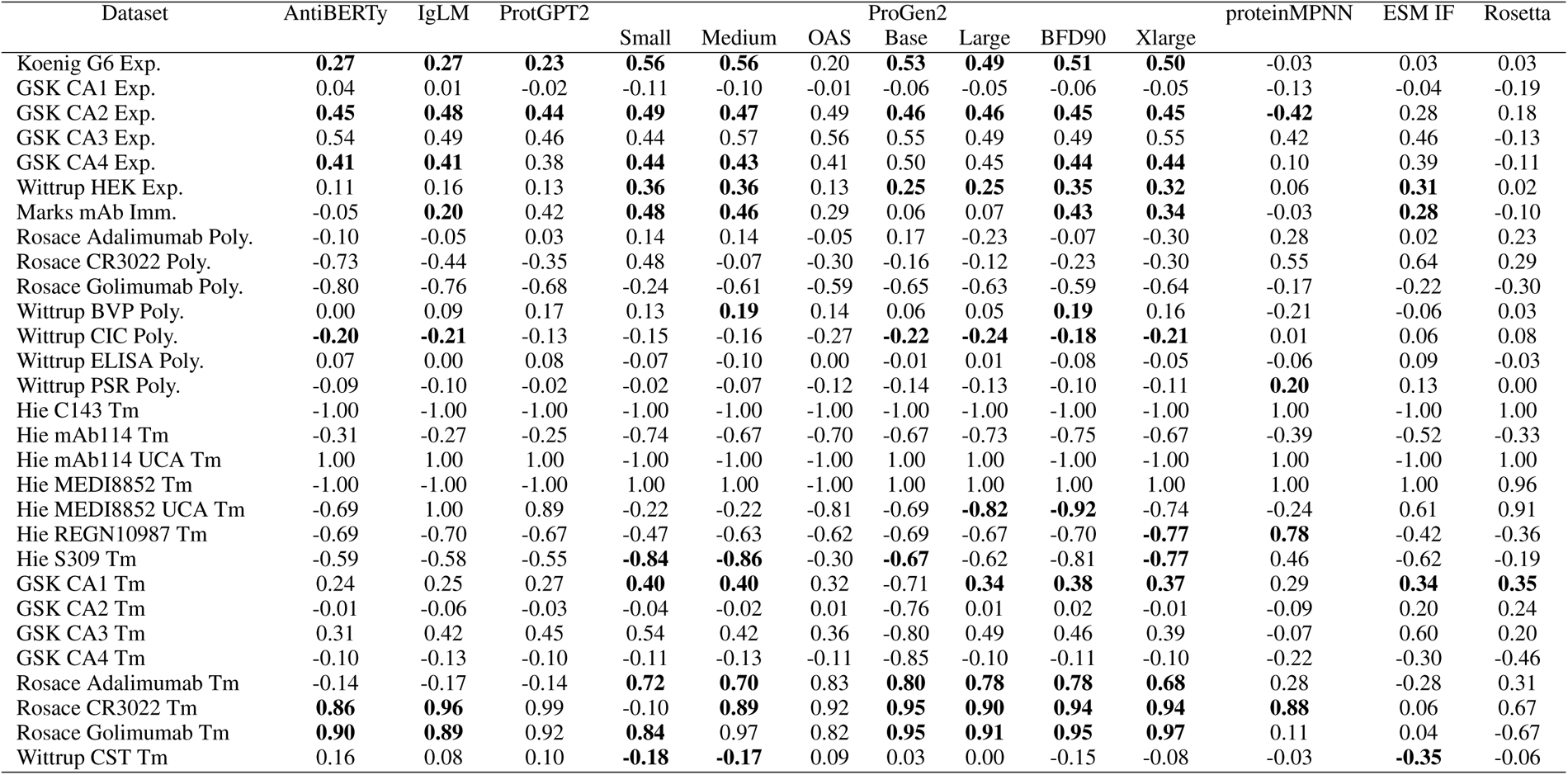
Summary of Pearson correlations (continued). Correlations with statistical significance (p < 0.05) are shown in bold.

**Table 3.** Summary of Spearman correlations. Correlations with statistical significance (p < 0.05) are shown in bold.

| Dataset | AntiBERTy | IgLM | ProTGPT2 | Small | Medium | OAS | ProGen2<br>Base | Large | BFD90 | Xlarge | proteinMPNN | ESM IF | Rosetta |
| --- | --- | --- | --- | --- | --- | --- | --- | --- | --- | --- | --- | --- | --- |
| Wittrup ACSINS Agg. | -0.16 | -0.16 | 0.01 | **-0.17** | **-0.18** | **-0.23** | -0.15 | **-0.17** | **-0.19** | **-0.18** | 0.09 | 0.14 | -0.03 |
| Wittrup CSI Agg. | -0.15 | **-0.20** | **-0.03** | **-0.17** | **-0.23** | -0.16 | **-0.19** | **-0.21** | **-0.25** | **-0.25** | 0.04 | 0.00 | -0.04 |
| Wittrup HIC Agg. | -0.05 | -0.03 | 0.14 | 0.02 | 0.00 | **-0.18** | -0.05 | -0.05 | -0.04 | -0.05 | **0.24** | **0.24** | 0.10 |
| Wittrup SAS Agg. | **0.20** | 0.07 | 0.24 | 0.04 | 0.02 | 0.01 | 0.16 | 0.13 | 0.02 | 0.04 | -0.05 | 0.12 | -0.01 |
| Wittrup SGAC Agg. | 0.11 | 0.10 | 0.27 | 0.07 | 0.09 | **0.26** | 0.09 | 0.10 | 0.10 | 0.10 | -0.06 | **-0.18** | 0.01 |
| Wittrup SMAC Agg. | -0.01 | -0.02 | 0.15 | -0.03 | -0.02 | **-0.20** | -0.02 | -0.03 | -0.04 | -0.05 | **0.27** | **0.18** | 0.06 |
| Hie C143 Kd | -0.06 | 0.01 | 0.01 | 0.28 | 0.18 | 0.04 | 0.36 | 0.11 | 0.00 | 0.03 | 0.14 | -0.01 | **-0.52** |
| Hie mAb114 Kd | -0.26 | -0.18 | -0.19 | -0.42 | -0.16 | 0.03 | -0.23 | -0.11 | -0.43 | -0.19 | -0.27 | -0.27 | -0.23 |
| Hie MED18852 Kd | 0.33 | 0.33 | 0.33 | **-0.61** | -0.32 | 0.11 | -0.05 | -0.13 | -0.08 | 0.05 | -0.02 | 0.12 | 0.07 |
| Hie MED18852 UCA Kd | **0.84** | **0.94** | 0.94 | 0.09 | 0.15 | **0.93** | **0.84** | **0.92** | **0.85** | **0.89** | -0.39 | 0.11 | **-0.60** |
| Hie REGN10987 Kd | 0.18 | 0.20 | 0.19 | 0.25 | 0.42 | 0.35 | 0.32 | 0.38 | 0.28 | 0.24 | 0.24 | 0.10 | -0.06 |
| Hie S309 Kd | **0.56** | **0.52** | **0.52** | **0.74** | **0.64** | 0.31 | **0.53** | **0.58** | **0.59** | **0.55** | 0.01 | 0.07 | -0.37 |
| GSK CA1 Kd | 0.11 | 0.05 | 0.05 | -0.04 | 0.02 | 0.04 | 0.03 | 0.05 | 0.01 | 0.03 | 0.06 | -0.12 | -0.05 |
| GSK CA2 Kd | **-0.83** | **-0.84** | **-0.84** | **-0.83** | **-0.82** | **-0.84** | **-0.85** | **-0.85** | **-0.83** | **-0.83** | 0.02 | **-0.63** | -0.08 |
| GSK CA3 Kd | 0.39 | 0.42 | 0.41 | 0.40 | 0.51 | 0.48 | 0.41 | 0.40 | 0.44 | 0.42 | 0.09 | 0.45 | -0.09 |
| GSK CA4 Kd | 0.35 | 0.37 | 0.37 | 0.36 | 0.46 | 0.44 | 0.36 | 0.36 | 0.39 | 0.37 | 0.05 | 0.41 | -0.14 |
| Koenig G6 Kd | **0.08** | **0.05** | **0.05** | **0.19** | **0.15** | **0.07** | **0.14** | **0.14** | **0.15** | **0.09** | 0.00 | 0.36 | -0.18 |
| Rosace Adalimumab Kd | -0.10 | -0.04 | -0.05 | 0.12 | 0.12 | -0.09 | 0.12 | -0.18 | 0.11 | -0.16 | -0.45 | 0.53 | -0.07 |
| Rosace CD3022 Kd | 0.09 | 0.03 | 0.03 | -0.77 | -0.49 | 0.03 | -0.26 | -0.26 | 0.03 | 0.03 | -0.31 | -0.37 | 0.14 |
| Rosace Golinumab Kd | 0.30 | 0.30 | 0.30 | 0.40 | 0.40 | 0.30 | 0.40 | 0.30 | 0.40 | 0.40 | 0.70 | -0.20 | -0.60 |
| Warsz D44 Kd | **0.45** | **0.42** | **0.41** | **0.26** | **0.30** | **0.25** | **0.39** | **0.40** | **0.33** | **0.35** | 0.01 | 0.05 | **0.07** |

**Table 4.** Summary of Spearman correlations (continued). Correlations with statistical significance (p < 0.05) are shown in bold.

| Dataset | AntiBERTy | IgLM | ProtGPT2 | Small | Medium | OAS | ProGen2<br>Base | Large | BFD90 | Xlarge | proteinMPNN | ESM IF | Rosetta |
| --- | --- | --- | --- | --- | --- | --- | --- | --- | --- | --- | --- | --- | --- |
| Koenig G6 Exp. | **0.24** | **0.21** | **0.18** | **0.45** | **0.40** | **0.18** | **0.39** | **0.38** | **0.38** | **0.33** | -0.03 | 0.01 | 0.02 |
| GSK CA1 Exp. | 0.12 | 0.08 | 0.04 | -0.06 | -0.09 | -0.01 | -0.02 | 0.00 | -0.06 | -0.03 | -0.13 | 0.14 | -0.11 |
| GSK CA2 Exp. | **0.60** | **0.62** | **0.58** | **0.63** | **0.60** | **0.62** | **0.60** | **0.62** | 0.59 | 0.59 | -0.40 | 0.36 | 0.14 |
| GSK CA3 Exp. | 0.55 | 0.58 | 0.54 | 0.49 | **0.75** | **0.63** | **0.62** | 0.55 | 0.54 | 0.58 | 0.55 | **0.64** | -0.08 |
| GSK CA4 Exp. | 0.39 | **0.41** | 0.37 | **0.42** | **0.48** | **0.42** | 0.57 | **0.44** | **0.46** | **0.45** | 0.07 | 0.28 | -0.06 |
| Wittrup HEK Exp. | 0.05 | 0.10 | 0.07 | **0.26** | **0.24** | **0.19** | **0.21** | **0.21** | **0.23** | **0.20** | 0.03 | **0.27** | -0.03 |
| Marks mAb Imm. | 0.13 | **0.24** | **0.27** | **0.32** | **0.33** | **0.31** | **0.15** | **0.15** | **0.32** | **0.27** | -0.02 | **0.14** | -0.01 |
| Rosace Adalimumab Poly. | 0.03 | 0.10 | 0.15 | 0.17 | 0.08 | 0.08 | 0.23 | -0.13 | 0.07 | -0.24 | 0.39 | -0.07 | 0.27 |
| Rosace CR3022 Poly. | 0.14 | 0.09 | 0.14 | 0.54 | 0.60 | 0.09 | 0.37 | 0.37 | 0.09 | 0.09 | 0.71 | **0.83** | 0.49 |
| Rosace Golimumab Poly. | -0.70 | -0.70 | -0.65 | -0.50 | -0.50 | -0.70 | -0.50 | -0.70 | -0.50 | -0.50 | -0.10 | -0.40 | -0.10 |
| Wittrup BVP Poly. | 0.02 | 0.13 | 0.18 | 0.10 | **0.17** | 0.12 | 0.07 | 0.08 | 0.18 | 0.18 | -0.15 | -0.04 | 0.00 |
| Wittrup CIC Poly. | **-0.19** | **-0.17** | -0.12 | -0.14 | **-0.18** | **-0.24** | **-0.18** | **-0.22** | **-0.20** | **-0.20** | 0.02 | 0.09 | 0.00 |
| Wittrup ELISA Poly. | -0.03 | -0.10 | -0.05 | -0.07 | -0.13 | -0.11 | -0.07 | -0.09 | -0.15 | -0.14 | -0.03 | 0.06 | 0.03 |
| Wittrup PSR Poly. | -0.11 | -0.14 | -0.09 | -0.09 | -0.14 | -0.08 | -0.16 | -0.15 | -0.17 | -0.16 | 0.15 | 0.10 | 0.00 |
| Hie C143 Tm | -1.00 | -1.00 | -1.00 | -1.00 | -1.00 | -1.00 | -1.00 | -1.00 | -1.00 | -1.00 | 1.00 | -1.00 | 1.00 |
| Hie mAb114 Tm | -0.11 | -0.11 | -0.10 | -0.68 | -0.71 | -0.54 | -0.57 | -0.71 | -0.71 | -0.68 | -0.29 | -0.61 | -0.18 |
| Hie mAb114 UCA Tm | 1.00 | 1.00 | 1.00 | -1.00 | -1.00 | -1.00 | 1.00 | 1.00 | -1.00 | -1.00 | 1.00 | -1.00 | 1.00 |
| Hie MED18852 Tm | -1.00 | -1.00 | -1.00 | 1.00 | 1.00 | -1.00 | 1.00 | 1.00 | 1.00 | 1.00 | 1.00 | 1.00 | 0.96 |
| Hie MED18852 UCA Tm | -0.77 | 1.00 | 1.00 | 0.14 | -0.14 | **-0.83** | **-0.83** | **-0.83** | -0.60 | **-0.83** | -0.31 | 0.77 | 0.91 |
| Hie REGN10987 Tm | **-0.71** | **-0.74** | -0.73 | -0.62 | **-0.83** | **-0.76** | **-0.76** | **-0.76** | **-0.81** | **-0.93** | **0.79** | -0.57 | -0.60 |
| Hie S309 Tm | **-0.71** | **-0.68** | **-0.68** | **-0.88** | **-0.83** | -0.55 | **-0.84** | **-0.68** | **-0.88** | **-0.77** | 0.35 | **-0.68** | 0.12 |
| GSK CA1 Tm | 0.16 | 0.21 | 0.21 | **0.40** | **0.38** | 0.23 | -0.89 | 0.29 | **0.38** | **0.35** | 0.20 | **0.35** | **0.36** |
| GSK CA2 Tm | -0.01 | -0.07 | -0.06 | -0.08 | -0.06 | -0.03 | -0.93 | -0.10 | -0.03 | -0.08 | -0.06 | 0.13 | 0.13 |
| GSK CA3 Tm | 0.24 | 0.29 | 0.29 | 0.45 | 0.26 | 0.26 | -0.98 | 0.45 | 0.45 | 0.29 | -0.21 | 0.31 | 0.29 |
| GSK CA4 Tm | -0.04 | -0.04 | -0.04 | -0.08 | -0.10 | -0.04 | -1.02 | -0.08 | -0.11 | -0.09 | **-0.41** | -0.28 | -0.07 |
| Rosace Adalimumab Tm | -0.08 | -0.09 | -0.08 | **0.62** | **0.55** | **0.79** | **0.68** | **0.64** | **0.64** | **0.56** | 0.21 | -0.24 | 0.33 |
| Rosace CR3022 Tm | **0.94** | **1.00** | 1.00 | -0.37 | **0.83** | **1.00** | **0.94** | **0.94** | **1.00** | **1.00** | **0.94** | -0.31 | 0.66 |
| Rosace Golimumab Tm | **0.90** | **0.90** | **0.90** | **1.00** | **1.00** | **0.90** | **1.00** | **0.90** | **1.00** | **1.00** | 0.30 | 0.40 | **-1.00** |
| Wittrup CST Tm | 0.14 | 0.05 | 0.06 | -0.14 | -0.12 | 0.05 | 0.02 | -0.01 | -0.11 | -0.07 | -0.02 | **-0.35** | -0.03 |

**Table 5.** Summary of Koendall tau correlations. Correlations with statistical significance (p < 0.05) are shown in bold.

| Dataset | AntiBERTy | IgLM | ProTGPT2 | Small | Medium | OAS | ProGen2<br>Base | Large | BFD90 | Xlarge | proteinMPNN | ESM IF | Rosetta |
| --- | --- | --- | --- | --- | --- | --- | --- | --- | --- | --- | --- | --- | --- |
| Wittrup ACSINS Agg. | -0.11 | -0.10 | **-0.10** | **-0.12** | **-0.12** | **-0.16** | -0.10 | -0.11 | **-0.13** | -0.11 | 0.06 | 0.10 | -0.03 |
| Wittrup CSI Agg. | -0.10 | **-0.14** | **-0.14** | **-0.12** | **-0.16** | -0.12 | **-0.13** | **-0.14** | **-0.18** | -0.17 | 0.03 | 0.00 | -0.03 |
| Wittrup HIC Agg. | -0.04 | -0.03 | -0.03 | 0.02 | 0.00 | **-0.12** | -0.04 | -0.05 | -0.03 | -0.04 | **0.17** | **0.16** | 0.07 |
| Wittrup SAS Agg. | **0.14** | 0.05 | 0.05 | 0.03 | 0.01 | 0.00 | 0.10 | 0.09 | 0.02 | 0.03 | -0.04 | 0.09 | -0.01 |
| Wittrup SGAC Agg. | 0.08 | 0.07 | 0.07 | 0.05 | 0.07 | **0.19** | 0.07 | 0.08 | 0.08 | 0.07 | -0.04 | **-0.13** | 0.01 |
| Wittrup SMAC Agg. | 0.00 | -0.01 | -0.01 | -0.01 | -0.01 | **-0.14** | -0.01 | -0.03 | -0.03 | -0.03 | **0.19** | **0.12** | 0.04 |
| Hie C143 Kd | -0.02 | 0.03 | 0.04 | 0.20 | 0.08 | 0.03 | 0.22 | 0.08 | -0.03 | 0.02 | 0.15 | 0.02 | -0.32 |
| Hie mAb114 Kd | -0.19 | -0.15 | -0.14 | -0.28 | -0.12 | 0.02 | -0.17 | -0.09 | -0.29 | -0.13 | -0.16 | -0.19 | -0.19 |
| Hie MED18852 Kd | 0.22 | 0.20 | 0.20 | **-0.41** | -0.26 | 0.05 | -0.10 | -0.22 | -0.18 | -0.03 | -0.01 | 0.07 | 0.03 |
| Hie MED18852 UCA Kd | **0.66** | **0.81** | 0.81 | 0.03 | 0.03 | **0.80** | **0.66** | **0.75** | **0.64** | **0.74** | -0.28 | 0.11 | **-0.47** |
| Hie REGN10987 Kd | 0.09 | 0.12 | 0.12 | 0.12 | 0.35 | 0.25 | 0.22 | 0.30 | 0.22 | 0.17 | 0.14 | 0.09 | -0.09 |
| Hie S309 Kd | **0.43** | **0.43** | **0.43** | **0.60** | **0.50** | 0.23 | **0.42** | **0.47** | **0.45** | **0.40** | 0.03 | 0.02 | -0.30 |
| GSK CA1 Kd | 0.09 | 0.03 | 0.03 | -0.05 | -0.01 | 0.04 | 0.00 | 0.03 | -0.03 | 0.00 | 0.04 | -0.11 | -0.04 |
| GSK CA2 Kd | **-0.66** | **-0.68** | **-0.68** | **-0.65** | **-0.64** | **-0.67** | **-0.69** | **-0.69** | **-0.65** | **-0.65** | 0.00 | **-0.52** | -0.07 |
| GSK CA3 Kd | 0.24 | 0.27 | 0.28 | 0.27 | 0.35 | 0.31 | 0.24 | 0.24 | 0.27 | 0.27 | 0.05 | 0.27 | -0.05 |
| GSK CA4 Kd | 0.19 | 0.23 | 0.23 | 0.23 | 0.30 | 0.26 | 0.19 | 0.19 | 0.23 | 0.23 | 0.01 | 0.23 | -0.10 |
| Koenig G6 Kd | **0.05** | **0.04** | **0.04** | **0.13** | **0.10** | **0.05** | **0.10** | **0.09** | **0.10** | **0.06** | **-0.04** | **0.18** | **-0.14** |
| Rosace Adalimumab Kd | -0.07 | -0.04 | -0.04 | 0.09 | 0.09 | -0.07 | 0.07 | -0.15 | 0.09 | -0.11 | -0.29 | 0.33 | -0.04 |
| Rosace CD3022 Kd | 0.07 | -0.07 | -0.06 | -0.60 | -0.33 | -0.07 | -0.20 | -0.20 | -0.07 | -0.07 | -0.20 | -0.33 | 0.33 |
| Rosace Golumumab Kd | 0.20 | 0.20 | 0.20 | 0.40 | 0.40 | 0.20 | 0.40 | 0.20 | 0.40 | 0.40 | 0.60 | -0.20 | -0.40 |
| Warsz D44 Kd | **0.31** | **0.28** | **0.29** | **0.18** | **0.21** | **0.17** | **0.27** | **0.27** | **0.23** | **0.24** | 0.01 | 0.02 | **0.05** |

**Table 6.** Summary of Kendall tau correlations (continued). Correlations with statistical significance (p < 0.05) are shown in bold.

| Dataset | AntiBERTy | IgLM | ProtGPT2 | Small | Medium | OAS | Base | Large | BFD90 | XLarge | proteinMPNN | ESM IF | Rosetta |
| --- | --- | --- | --- | --- | --- | --- | --- | --- | --- | --- | --- | --- | --- |
| Koenig G6 Exp. | **0.16** | **0.14** | **0.15** | **0.31** | **0.28** | **0.12** | **0.27** | **0.25** | **0.26** | **0.22** | -0.04 | -0.03 | 0.00 |
| GSK CA1 Exp. | 0.09 | 0.06 | 0.06 | -0.04 | -0.06 | -0.02 | 0.00 | 0.01 | -0.02 | 0.00 | -0.09 | 0.11 | -0.07 |
| GSK CA2 Exp. | **0.42** | **0.43** | **0.44** | **0.44** | **0.39** | **0.40** | **0.38** | **0.43** | **0.39** | **0.39** | **-0.29** | 0.24 | 0.13 |
| GSK CA3 Exp. | 0.42 | 0.45 | 0.46 | 0.38 | **0.60** | **0.49** | 0.49 | 0.42 | 0.38 | 0.45 | 0.27 | **0.49** | -0.09 |
| GSK CA4 Exp. | 0.25 | 0.26 | 0.26 | 0.28 | **0.31** | 0.28 | 0.45 | 0.28 | **0.31** | **0.30** | 0.04 | 0.17 | -0.02 |
| Wittrup HEK Exp. | 0.04 | 0.07 | 0.07 | **0.18** | **0.16** | **0.12** | **0.14** | **0.14** | **0.15** | **0.13** | -0.01 | **0.18** | -0.02 |
| Marks mAb Imm. | 0.09 | **0.16** | **0.17** | **0.23** | **0.24** | **0.22** | **0.10** | **0.10** | **0.23** | **0.19** | -0.02 | **0.09** | -0.01 |
| Rosace Adalimumab Poly. | 0.03 | 0.05 | 0.06 | 0.14 | 0.05 | 0.08 | 0.16 | -0.10 | 0.05 | -0.14 | 0.27 | -0.03 | 0.19 |
| Rosace CR3022 Poly. | 0.33 | 0.20 | 0.20 | 0.47 | 0.47 | 0.20 | 0.33 | 0.33 | 0.20 | 0.20 | 0.47 | 0.73 | 0.33 |
| Rosace Golimumab Poly. | -0.60 | -0.60 | -0.60 | -0.40 | -0.40 | -0.60 | -0.40 | -0.60 | -0.40 | -0.40 | 0.00 | -0.40 | 0.00 |
| Wittrup BVP Poly. | 0.01 | 0.08 | 0.08 | 0.07 | 0.11 | 0.08 | 0.05 | 0.05 | 0.11 | **0.11** | -0.05 | -0.03 | 0.00 |
| Wittrup CIC Poly. | **-0.14** | **-0.12** | -0.11 | **-0.10** | **-0.13** | **-0.16** | **-0.13** | **-0.16** | **-0.13** | **-0.14** | 0.02 | 0.06 | -0.01 |
| Wittrup ELISA Poly. | -0.02 | -0.07 | -0.06 | -0.05 | -0.09 | -0.08 | -0.04 | -0.05 | -0.09 | -0.09 | -0.02 | 0.04 | 0.02 |
| Wittrup PSR Poly. | -0.07 | -0.10 | -0.10 | -0.07 | -0.10 | -0.06 | -0.11 | -0.10 | -0.12 | -0.12 | 0.11 | 0.07 | 0.00 |
| Hie C143 Tm | -1.00 | -1.00 | -1.00 | -1.00 | -1.00 | -1.00 | -1.00 | -1.00 | -1.00 | -1.00 | 1.00 | -1.00 | 1.00 |
| Hie mAb114 Tm | -0.05 | -0.05 | -0.04 | -0.52 | -0.62 | -0.43 | -0.43 | -0.62 | -0.62 | -0.52 | -0.14 | -0.52 | -0.05 |
| Hie mAb114 UCA Tm | 1.00 | 1.00 | 1.00 | -1.00 | -1.00 | -1.00 | 1.00 | 1.00 | -1.00 | -1.00 | 1.00 | -1.00 | 1.00 |
| Hie MED18852 Tm | -1.00 | -1.00 | -1.00 | 1.00 | 1.00 | -1.00 | 1.00 | 1.00 | 1.00 | 1.00 | 1.00 | 1.00 | 0.96 |
| Hie MED18852 UCA Tm | -0.60 | 1.00 | 1.00 | 0.07 | -0.07 | -0.73 | -0.73 | -0.73 | -0.47 | -0.73 | -0.20 | 0.60 | 0.91 |
| Hie REGN10987 Tm | -0.57 | **-0.64** | -0.64 | -0.43 | **-0.71** | -0.57 | -0.57 | -0.57 | **-0.64** | **-0.86** | **0.64** | -0.50 | -0.43 |
| Hie S309 Tm | **-0.56** | **-0.56** | **-0.55** | **-0.73** | **-0.69** | -0.42 | **-0.64** | -0.47 | **-0.73** | **-0.64** | 0.20 | **-0.51** | 0.16 |
| GSK CA1 Tm | 0.11 | 0.15 | 0.15 | **0.27** | **0.26** | 0.17 | -0.69 | 0.19 | **0.26** | 0.23 | 0.13 | **0.26** | **0.25** |
| GSK CA2 Tm | -0.01 | -0.04 | -0.04 | -0.04 | -0.04 | -0.01 | -0.73 | -0.07 | -0.01 | -0.04 | -0.04 | 0.09 | 0.10 |
| GSK CA3 Tm | 0.29 | 0.36 | 0.36 | 0.43 | 0.29 | 0.29 | -0.78 | 0.43 | 0.43 | 0.36 | -0.07 | 0.21 | 0.14 |
| GSK CA4 Tm | -0.03 | -0.03 | -0.03 | -0.06 | -0.07 | -0.04 | -0.82 | -0.05 | -0.08 | -0.07 | -0.30 | -0.23 | -0.04 |
| Rosace Adalimumab Tm | -0.07 | -0.08 | -0.07 | **0.42** | **0.42** | **0.62** | **0.53** | **0.53** | **0.51** | **0.40** | 0.18 | -0.18 | 0.27 |
| Rosace CR3022 Tm | **0.87** | **1.00** | 1.00 | -0.33 | 0.73 | **1.00** | **0.87** | **0.87** | **1.00** | **1.00** | **0.87** | -0.33 | 0.47 |
| Rosace Golimumab Tm | 0.80 | 0.80 | 0.80 | **1.00** | **1.00** | 0.80 | **1.00** | 0.80 | **1.00** | **1.00** | 0.20 | 0.40 | -1.00 |
| Wittrup CST Tm | 0.09 | 0.04 | 0.04 | -0.10 | -0.08 | 0.03 | 0.01 | -0.01 | -0.07 | -0.05 | -0.01 | **-0.24** | -0.02 |

